# Dynamic modelling and analysis of autophagy in the clearance of aggregated α-synuclein in Parkinson’s disease

**DOI:** 10.1101/2022.08.26.505373

**Authors:** Bojie Yang, Zhuoqin Yang, Heng Liu

## Abstract

The widely-accepted hallmark pathology of Parkinson’s disease (PD) is the presence of Lewy bodies (LB) with characteristic abnormal aggregated α-synuclein (αSyn). Growing physiological evidences suggest that there is a pivotal role for the autophagy-lysosome pathway in the clearance of misfolded αSyn (αSyn*) for maintaining homeostasis and neural cell function. In this work, we establish a new mathematical model for αSyn* degradation through the autophagy pathway. The qualitative simulations discover the tri-stability phenomena and dynamical behaviors of αSyn*, i.e., the coexistence of three stable steady states, in which the lower, medium and upper steady states correspond to the healthy, critical and diseased stages of pathological mechanism of PD, respectively. Diverse analyses on codimension-1 and -2 bifurcations suggest that autophagy can control the switches among the stable steady states for the aggregation of αSyn*. It is also found that the double negative crosstalk feedback between autophagy and apoptosis is important to the robustness of tri-stability of αSyn* for this biodynamic system. Our novel results may be valuable for making further therapeutic strategies in prevention and treatment for PD.

**Author summary:** αSyn* is one of the most and primary remarkable targets for the universal neurodegenerative disease of PD. Efficient clearance mechanism of autophagy, contains a lot of molecular components to maintain the cellular homeostasis and cell renewal, could be the possible therapy for PD. Understand the complexity of autophagy in degrading αSyn* requires the integration of both theoretical and experimental points of view. Here, we have proposed a novel mathematical model that analyses the temporal and dynamic behaviors of the biosystem for autophagy degrades αSyn* to access PD states. The model explains that the tri-stability is of particularly relevance to this biosystem that switch among the healthy, critical and disease states, and captures that the critical intermedium state exits for preventing the system transform healthy to disease state directly, and further illustrates that the molecular signaling feedback loops of autophagy may be important for the robustness of tri-stability. Our work deepens the researches of PD by uncovering the important buffering medium state and providing promising potential therapeutic insights.

## 1 Introduction

Parkinson’s disease (PD), widely known as a severe progressive neurodegenerative disorder, has affected millions of people worldwide [1–3]. Even though the intraneuronal accumulation of α-synuclein (αSyn) is critical for PD development, it is expected to further uncover the related molecular mechanisms that may trigger the disease [4–6].

Aggregated αSyn is a major component of Lewy bodies (LB), the pathological hallmark of PD [7–10]. Several mathematical approaches to understand the kinetics of αSyn aggregation relevance to PD have been analyzed by former researches [11, 12]. The common characteristic of PD is mitochondria dysfunction, caused by several mechanisms, e.g., aggregation of misfolded αSyn (αSyn*), increasement of reactive oxygen species (ROS), homeostasis of calcium and impairments of mitochondria dynamics [13–15]. The various molecules and cellular processes mentioned above include a vicious circle between αSyn* and ROS, which has been used by Finnerty *et al*. to predict a systematic and irreversible switch to deadly high levels of ROS after enough exposure to risk factors associated with PD [16]. Moreover, the dynamic behaviors of the circulations between αSyn* and ROS have been surveyed by Cloutier *et al*. by a mathematical model [17, 18], in which the central motif highlights the double positive feedback loop between αSyn* and ROS with the input signals of S_1_ (*i* = 1 – 4) to analyze the pathology mechanisms of PD.

One possible method to cure PD is to promote the degradation of αSyn* in two important ways: the ubiquitin-proteasome system (UPS) and autophagy lysosomal pathway (ALP) [19–23]. Recent evidence suggests that the clearance of autophagy might play a major role in degrading αSyn* than proteasome [24–27]. Autophagy (i.e., “self-eating” process) is an essential metabolic pathway that degrades proteins and organelles in lysosome to preserve the cellular homeostasis [28–32]. Under mild stress conditions, autophagy acts as a survival mechanism while severe stress may switch on programmed cell death of apoptosis (i.e., “self-killing” process) [33–36]. Besides, the unfolded or misfolded proteins in the endoplasmic reticulum (ER), e.g., αSyn*, trigger ER stress response called unfolded protein response (UPR) and further activate the crosstalk between autophagy and apoptosis [37]. The comprehensive process, depicted by Holczer *et al*. through a mathematical model with ER stress sensors (ERS), directly activates the autophagy and apoptosis inducers [38]. Moreover, in the mathematical models proposed by Kapuy *et al*., Beclin1 and Caspases work as the main autophagy and apoptosis inducers, respectively [39].

The interactions between αSyn* and ROS reveal the basic mechanism of the aggregation of αSyn*, and the crosstalk between autophagy and apoptosis illustrates the removal regulation of abnormal aggregated proteins in vivo. The study of these processes is on the bases of both physiological and molecular levels. The dynamics of regulation of αSyn* degraded through the autophagy pathway via various protein regulators is still unknown [40, 41]. It is necessary to explore the essential complex interactions among molecular components in order to understand the clearance of autophagy in degrading αSyn*. In other words, it is needed to make deeper investigation on the functions and correlations of individual elements and the complexity of dynamic behaviors based on molecular biology and physiology [42].

In this paper we aim to explore the degradation of aggregated αSyn* through autophagy from a biophysical viewpoint. From this perspective, we establish a novel mathematical model of autophagy for the degradation of abnormal aggregated αSyn*. In our model, the aggregated abnormal αSyn* promoted by ROS activates ERS and then results in Beclin1-induced autophagy and Caspases-induced apoptosis of neuron cells. Bifurcation analyses of complex dynamic properties and the pathological phenomena of multi-stability of αSyn* are explored. Further, the aggregation of αSyn* under different initial conditions and the robustness of tri-stability of αSyn* are also concerned in this study. The obtained results may shed new light to understand the complexity of dynamic processes and mechanisms in the pathology and therapy of PD.

## 2 Model

### Model of aggregated αSyn* degraded by autophagy

Our mathematical model contains three processes, i.e., aggregation (orange block), autophagy (green block) and apoptosis (blue block) (Fig 1). Aggregated αSyn* is digested through Beclin1-dependent autophagy, however, for too large amount of αSyn* the system turns on Caspases-dependent apoptosis instead of autophagy. A positive feedback loop for the aggregation is formed between ROS and αSyn*. The internal or external oxidative stresses denoted by *S*_1_ promote ROS, while the age-related anti-oxidative mechanism, for instance, the decreased energy metabolism marked by *S*_2_ degrades ROS. And the influence of genetic damage or mutation is characterized by *S*_3_ to promote the aggregation of αSyn*. Furthermore, αSyn* activates ERS (the upper red arrow), which activates both Beclin1-dependent autophagy and Caspases-dependent apoptosis via a double negative feedback loop simultaneously. Especially, Beclin1-dependent autophagy acts as the major protein clearance mechanism for degrading αSyn* (the lower red arrow).

**FIG. 1.**
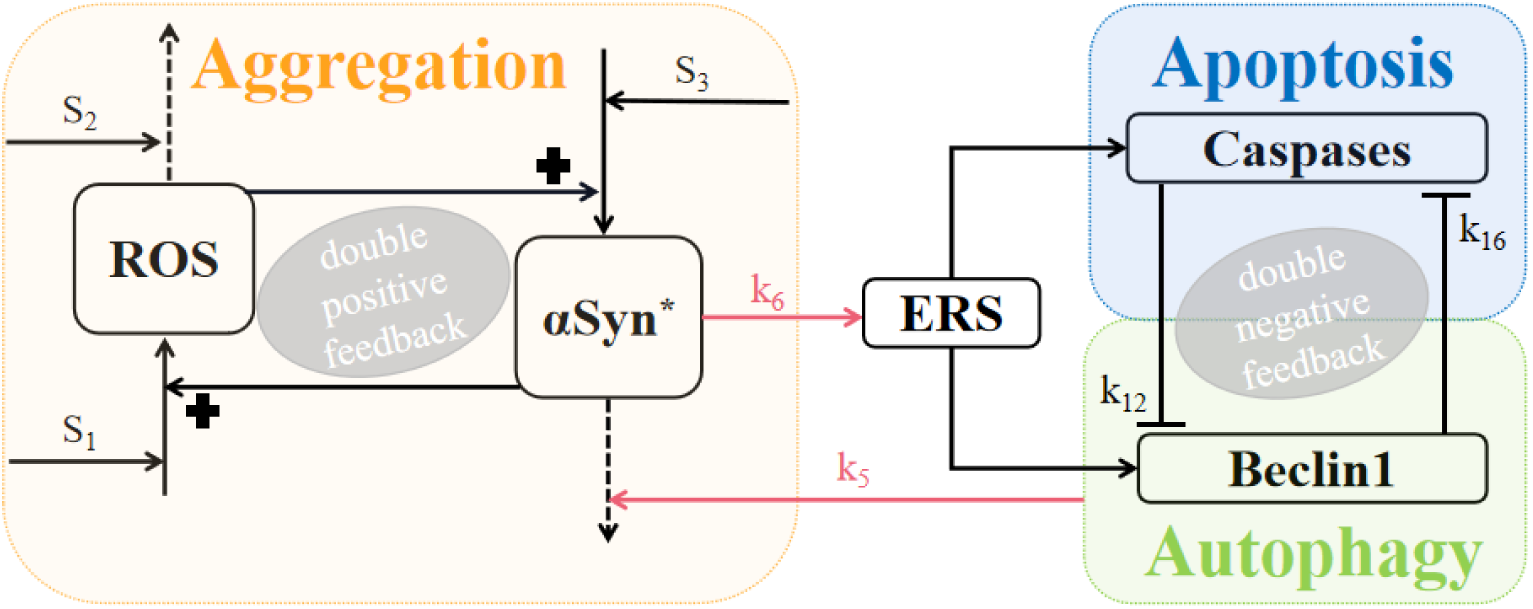
Schematic of the network model of aggregation, autophagy and apoptosis. The promotion, degradation and inhibition are denoted by solid lines with arrowheads, dotted lines with arrowheads and blocked end lines, respectively.

Based on the biochemical interactions (Fig 1), in this model we consider five components: ROS ([*R*]), αSyn* ([α*S*]), ERS ([*E*]), Beclin1 ([*B*]) and Caspases ([*C*]). The dynamics of the module is described by five nonlinear ordinary differential equations in our mathematical model. The description of the parameters is given and their values for computation are based on biological experiments (Table 1). The numerical simulations and bifurcation analysis for the system are performed by means of XPP, and the numerical solutions of the ordinary differential equations are obtained by the Runge-Kutta method in very high precision.

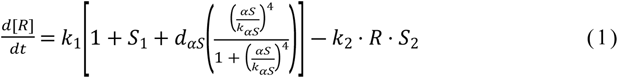

**Table 1.** Parameters of functions and their descriptions.

| Parameter | Significance | Value |
| --- | --- | --- |
| $S_1$ | Internal and external oxidative stresses | 2 |
| $S_2$ | Age-related anti-oxidative mechanisms | 1 |
| $S_3$ | Influence of genetic damage/mutation | 1 |
| $d_{\alpha S}$ | Dimensionless | 15 |
| $k_{\alpha S}$ | Hill constant | 8.5 |
| $k_1$ | Generation rate of ROS | 0.72 |
| $k_2$ | Removal time of ROS | 0.72 |
| $k_3$ | Generation time of $\alpha\text{Syn}^*$ | 0.7 |
| $k_4$ | Removal time of $\alpha\text{Syn}^*$ | 0.7 |
| $k_5$ | Beclin1-dependent rate of $\alpha\text{Syn}^*$ degradation | 2.7 |
| $k_6$ | $\alpha\text{Syn}^*$-dependent rate of ERS activation | 1 |
| $k_7$ | Basal rate of ERS activation | 0.5 |
| $k_8$ | Basal rate of ERS inactivation | 1.6 |
| $k_9$ | Basal rate of Beclin1 activation | 2 |
| $k_{10}$ | ERS-dependent rate of Beclin1 activation | 4 |
| $k_{11}$ | Basal rate of Beclin1 inactivation | 3.6 |
| $k_{12}$ | Caspases-dependent rate of Beclin1 inactivation | 10 |
| $k_{13}$ | Basal rate of Caspases inactivation | 1.8 |
| $k_{14}$ | ERS-dependent rate of Caspases activation | 2.2 |
| $k_{15}$ | Basal rate of Caspases inactivation | 2 |
| $k_{16}$ | Beclin1-dependent rate of Caspases inactivation | 4.5 |
| $Jbe$ | Beclin1 Michaelis constant | 1 |
| $Jca$ | Caspases Michaelis constant | 0.04 |
| $ET$ | Total level of ERS | 2 |
| $mT$ | Total level of mTOR | 1 |
| $BT$ | Total level of Beclin1 | 1 |
| $CT$ | Total level of Caspases | 1 |

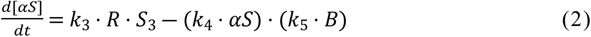

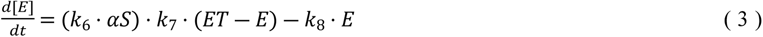

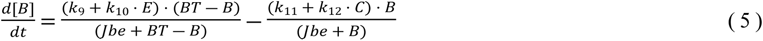

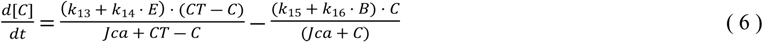

## 3 Results

### Bifurcation analysis and temporal behaviors of αSyn*, Caspases and Beclin1 under *S*_1_

More and more researchers focus on the mitochondrial dysfunction in PD related to the aggregation of αSyn* [43–46]. Mitochondria dysfunction is not only one of the important reasons to increase the stress *S*_1_ but also the crucial place to generate ROS, which accelerates the promotion of αSyn* within the double positive feedback loop. Therefore, we strive to investigate the dynamic behaviors of αSyn* under *S*_1_ in specialty, which may provide new insight from biodynamic aspect to discuss mitochondria dysfunction affects αSyn*.

To understand how *S*_1_ influences the dynamic bifurcation and temporal behaviors of αSyn*, Caspases and Beclin1, we consider the codimension-one bifurcation diagrams and time evolutions obtained by numerical simulations. The multiple steady states of αSyn* (Fig 2A1), Caspases (Fig 2B1) and Beclin1 (Fig 2C1) are given as the functions of S_1_, respectively. The fold bifurcation points *F_i_* (*i* = 1,2,…,5) evoke the coexistence of multiple stable steady states (denoted by red, yellow and green solid lines) and unstable ones (denoted by black dotted line). From these results we can observe the phenomena of tri-stability (yellow), bistability (pink) and mono-stability (blue), characterized by the number of the coexisting stable steady states at fixed *S*_1_. There exists tri-stability in the bifurcation diagrams of αSyn* (Fig 2A1), Caspases (Fig 2B1) and Beclin1 (Fig 2C1) within the range [0.035,1.365] of *S*_1_, respectively. It is worth noting that since proteins with nonnegative values are of biological sense in practice, we mainly consider the region enclosed by gray dotted lines on (*S*_1_,*y*)-plane, where *y* represents αSyn*, Caspases or Beclin1, respectively.

**FIG. 2.**
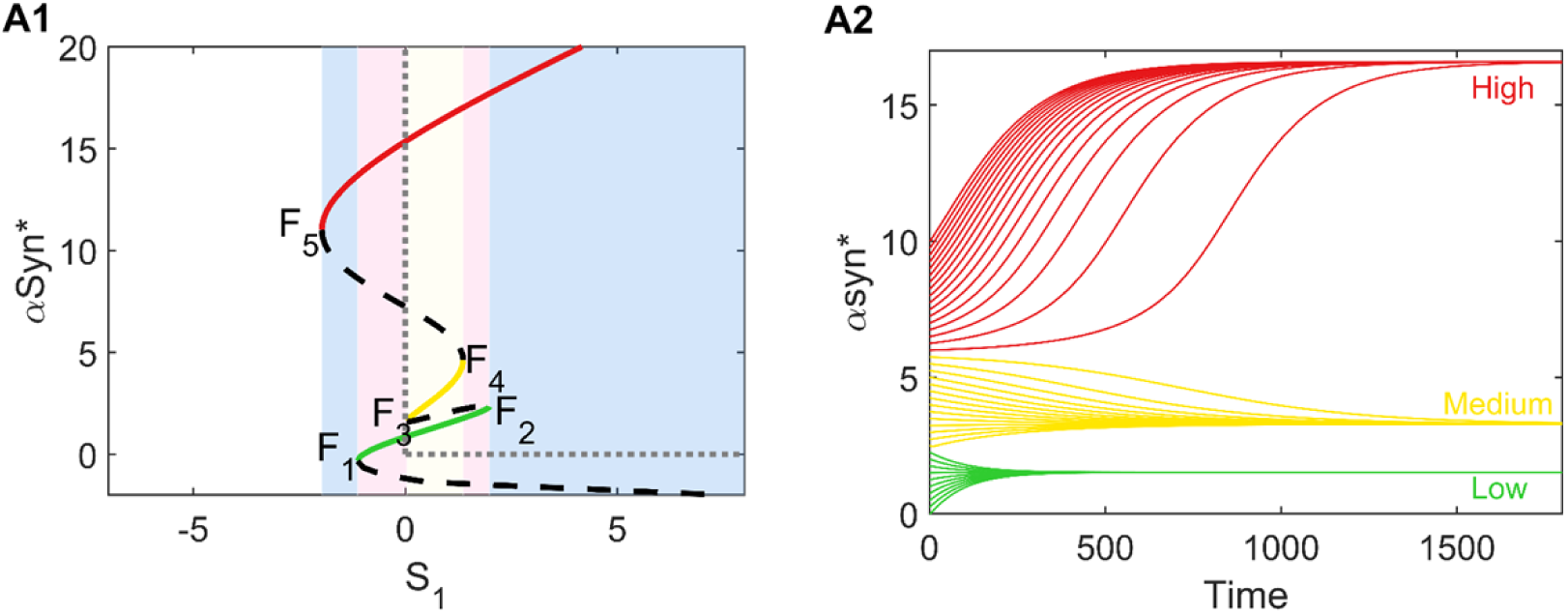
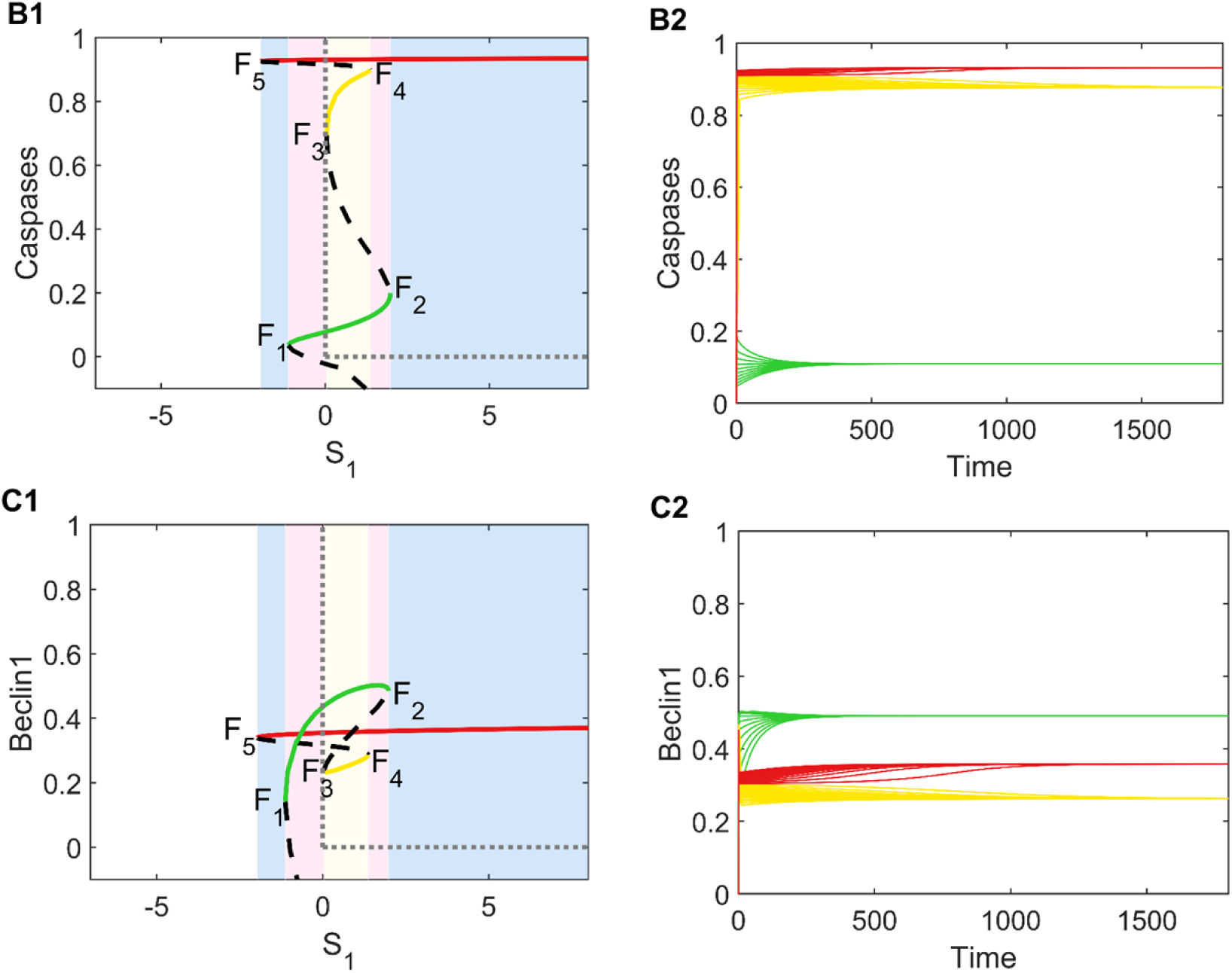
Bifurcation diagrams and temporal dynamics of αSyn*, Caspases and Beclin1. In the bifurcation diagrams of αSyn* (A1), Caspases (B1) and Beclin1 (C1) with respect to the parameter *S*_1_ (the input signal), *F_i_* (*i* = 1, …, 5) are fold bifurcation points, and the stable and unstable steady states are represented by colorful solid and black dotted lines, respectively. For *S*_1_ = 1, the time series of αSyn*(A2), Caspases (B2) and Beclin1 (C2) from different initial conditions (green, red and yellow) reach their stable steady states, respectively.

Taking account of that the aggregation of αSyn* is a main disease cause of PD, the pathological influence of the dynamical properties of tri-stability of αSyn* is concerned here. There are three (i.e., lower, medium and upper) stable steady states and only *F_i_* (*i* = 2, 3, 4) among the fold bifurcation points in the right half part of (*S*_1_,α*S*yn*)-plane. Obviously, the lower stable steady state with low concentration of αSyn* corresponds to the healthy stage of pathological mechanism of PD, and the upper stable steady state with high concentration of αSyn* corresponds to the diseased stage of pathological mechanism of PD. It is also can be seen (Fig 2A1) that the lower stable steady state may switch to the upper one directly via the fold bifurcation at *F*_2_, that is, it is possible to switch from the healthy stage to the diseased stage of PD directly due to the variation of *S*_1_. Now we further consider the role of the medium stable steady state of αSyn*. Because both of the fold bifurcation points *F*_3_ and *F*_4_ are in the positive half axis of (*S*_1_,α*S*yn*)-plane, the medium stable steady state may switch to either the lower one (i.e., the healthy stage of PD) via the fold bifurcation at *F*_3_ or the upper one (i.e., the diseased stage of PD) via the fold bifurcation at *F*_4_ (i.e., the reversibility of the bistable switch via fold bifurcation). This means that the medium stable steady state corresponds to the critical stage of PD, in which the disease situation may be improved or become worse depending on the variation of *S*_1_. On the contrary, the upper stable steady state cannot switch to other stable steady states via the fold bifurcation at *F*_5_ in the right half part of (*S*_1_,α*S*yn*)-plane (i.e., the irreversibility of the bistable switch via fold bifurcation). Hence, it is impossible to improve the situation of the diseased stage of PD by the variation of *S*_1_.

Now we consider the influence of initial values of αSyn*. Here, the time courses of the concentrations of αSyn* (Fig 2A2), Caspases (Fig 2B2) and Beclin1 (Fig 2C2) initiated from different initial values for *S*_1_ =1, respectively. From the regimes of tri-stability (Fig 2 A1, B1 and C1), it should be noticed that the lower, medium and upper stable steady states of both αSyn* and Caspases exactly correspond to the upper, lower and medium states of Beclin1, respectively. Firstly, small initial values of the concentration of αSyn* result in the state with low concentrations of αSyn* and Caspases but high concentration of Beclin1. This fact means that Beclin1-induced autophagy can degrade abundant αSyn* to keep low concentration of αSyn* for the healthy stage of PD. Secondly, medium initial values of the concentration of αSyn* cause the state with medium concentrations of αSyn* and Caspases but low concentration of Beclin1. This means that Caspases-induced apoptosis comes into effect for the critical stage of PD. Finally, large initial values of the concentration of αSyn* lead to the state with high concentration of αSyn* and Caspases but medium concentration of Beclin1. This means that Caspases-induced apoptosis plays a dominant role in the diseased stage of PD.

In conclusion, all of the above confirm the dynamical properties of tri-stability of αSyn*, i.e., the coexistence of three stable steady states of αSyn*, in which the lower, medium and upper states correspond to the healthy, critical and diseased stages of PD, respectively. The three stages of PD can be characterized by the level of Beclin1-induced autophagy or Caspases-induced apoptosis, which primarily works during the degradation of αSyn* and is governed by the temporal behaviors of αSyn*, Caspases and Beclin1 dependent on initial values. It is found that both the healthy and critical stages may transit to the diseased stage of PD with the increase of *S*_1_. Moreover, PD is possible to be cured by proper medical measures in the critical stage, although the treatment for PD in the diseased stage is still a crucial and difficult problem.

### Multistable steady states of αSyn* under the control of the parameters *k*_5_ and *k*_6_

Now we consider the aggregated αSyn* degraded mainly through the autophagy pathway in the mathematical model (1) - (6). Here, we focus on the key parameters *k*_5_ (Beclin1-dependent rate of αSyn* degradation) and *k*_6_ (αSyn*-dependent rate of ERS activation) connecting the two modules of aggregation and autophagy.

In order to dig out the dynamic impacts of autophagy on degrading αSyn*, we consider the bifurcation diagrams of αSyn* with respect to *S*_1_ for different values of *k*_5_ (Fig 3A1-A6). The ranges of tri-stability (yellow), bistability (pink) and mono-stability (blue) are presented. It is seen that tri-stability only appears for moderate values of *k*_5_ (Fig 3A3 and A4). A remarkable finding is that the bistable switches on the bifurcation curves alter from irreversible to reversible as the bifurcation curves move from the left to right with the increase of *k*_5_ in (*S*_1_,α*S*yn*)-plane. For large *k*_5_, the reversibility of bistable switch between the medium and upper stable steady states (i.e., the critical and the disease stages) pulls the high concentration of αSyn* down largely, which implies that autophagy-independent pathway plays an important role in degrading the aggregated αSyn* [27, 40, 47, 48].

**FIG. 3.**
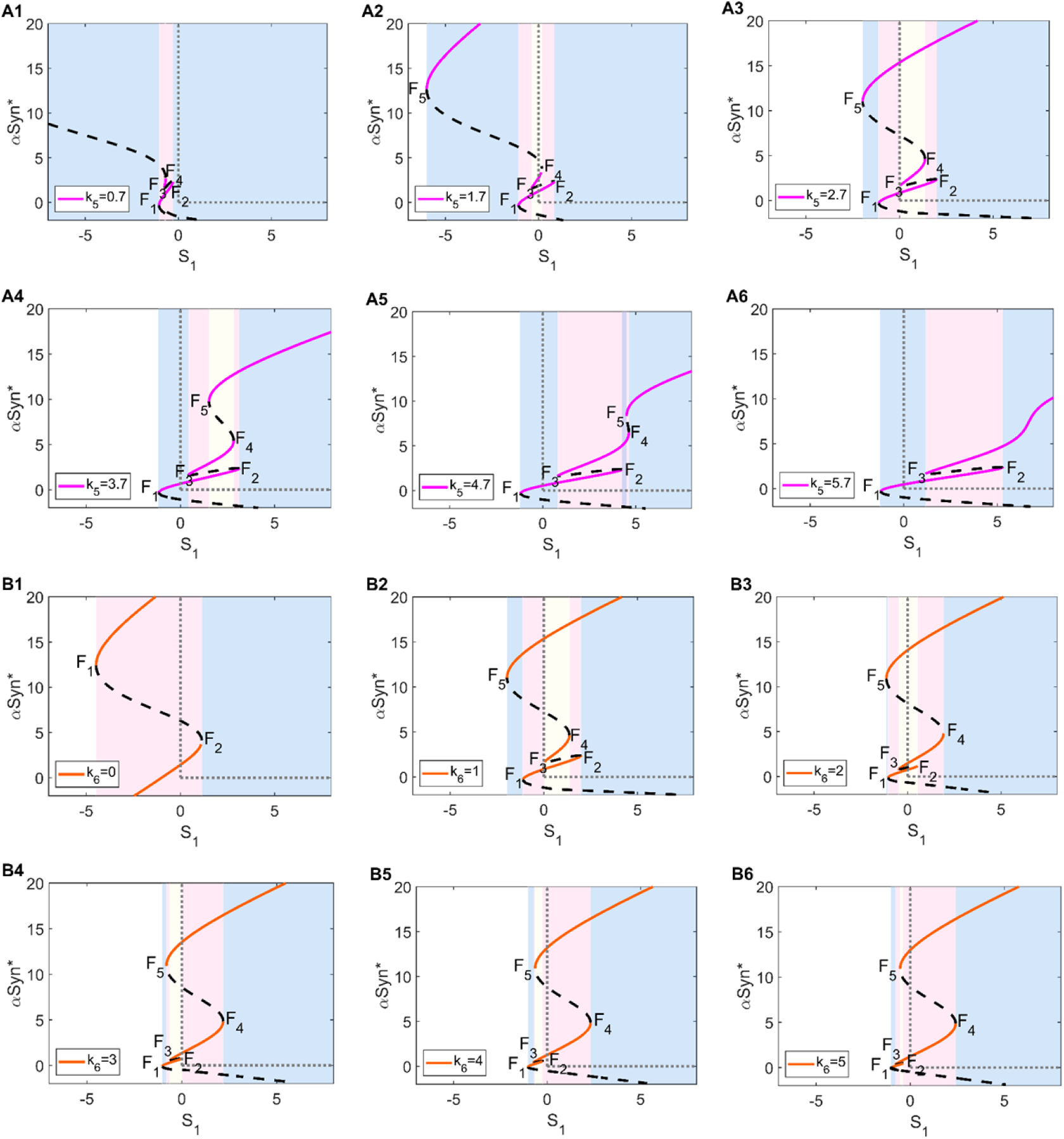
Bifurcation diagrams of αSyn* with respect to *S*_1_ under different values of *k*_5_ (A1 to A6) and *k*_6_ (B1 to B6).

In addition, we turn to the bifurcation diagrams of αSyn* with respect to *S*_1_ for different values of *k*_6_ (Fig 3B1-B6). The bifurcation curves shift slightly from left to right, and tri-stability also exists for moderate values of *k*_6_ (Fig 3B2 and B3). However, when *k*_6_ is large enough, the range of the lower stable steady state (i.e., the healthy stage) gradually shrinks and eventually falls onto the left half part of (*S*_1_,α*S*yn*)-plane, while the medium and upper stable steady states (i.e., the critical and diseased stages) with irreversible bistable switches coexist in the right half part of (*S*_1_, α*S*yn*)-plane (Fig 3B5 and B6). These results indicate that the parameter *k*_6_ hardly controls the high level of αSyn* for large stress *S*_1_ in PD treatment.

We further investigate all possible stable steady states in (*S*_1_,*k*_5_)- and (*S*_1_,*k*_6_) – planes to consider the regulations of *k*_5_ and *k*_6_ on multi-stability by the codimension-two bifurcation diagrams (Fig 4A and 4B), respectively. There are the regions of tri-stability (T), bistability (B) and mono-stability (M) divided by the fold bifurcation curves *f_i_* (*i* = 1,…, 5) (Fig 4) from the saddle-node bifurcation points *F_i_* (*i* = 1,…, 5) (Fig 3). Three pairs of fold bifurcation curves *f*_1_ and *f*_2_, *f*_3_ and *f*_4_, *f*_4_ and *f*_5_ intersect at the cusp bifurcation points CP3, CP2 and CP1 in (*S*_1_,*k*_5_)-plane, respectively, while the fold bifurcation curves *f*_3_ and *f*_4_ intersect at the cusp bifurcation point CP1 in (*S*_1_,*k*_6_)-plane. In addition, parallel to the codimension-one bifurcation diagrams (Fig 3), we add the pink and orange thin lines at the corresponding six values of *k*_5_ and *k*_6_ (Fig 3A and 3B), respectively. Through the codimension-two bifurcation diagrams in the (*S*_1_,*k*_5_)- and (*S*_1_,*k*_6_)-planes, we can observe the multi-stability in the regions M, B and T with coexistence of steady states more comprehensively, especially the occurrence of tri-stability in the region T, in which the healthy state may transit to the diseased state indirectly through buffering via the critical state as an alert warning for timely treatments to prevent PD.

**FIG. 4.**
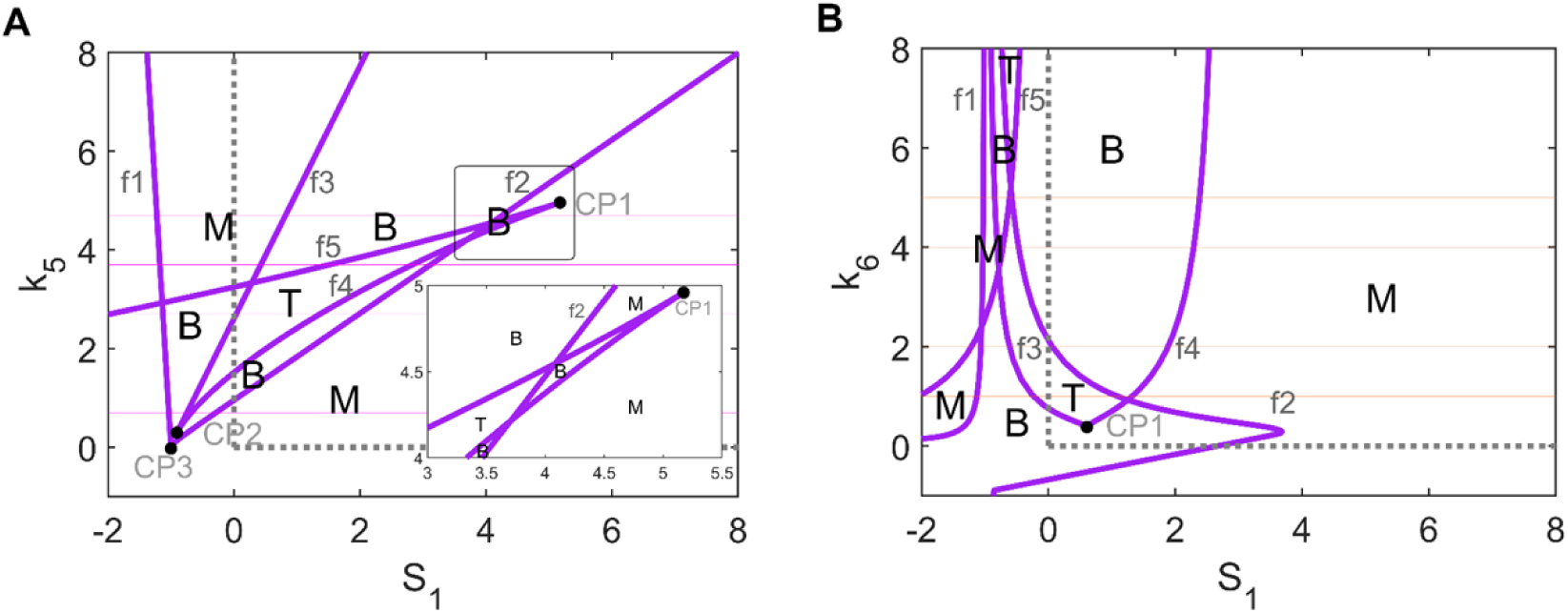
Codimension-two bifurcation diagram with respect to *S*_1_ and *k*_5_(A) (or *k*_6_ (B)). T, B and M denote the regions of tri-stability, bistability and mono-stability, respectively. Three pairs of fold bifurcation curves *f*_1_ and *f*_2_, *f*_3_ and *f*_4_, *f*_4_ and *f*_5_ (purple solid curves) intersect at the cusp bifurcation points CP3, CP2 and CP1 in (*S*_1_,*k*_5_)-plane, respectively, and *f*_3_ and *f*_4_ intersect at the cusp bifurcation point CP1 in (*S*_1_,*k*_6_)-plane. The pink and orange thin lines represent the six values of *k*_5_ and *k*_6_ from Fig 3A and 3B, respectively.

### The aggregation of αSyn* under different initial conditions

An important dynamic characteristic of tri-stability is that the system can switch among the stable steady states, which correspond to different stages in the pathological mechanism of PD. For a fixed set of parameters, initial values play a key role in deciding their final states as shown in the time series of αSyn*, Caspases and Beclin1 (Fig 2). From a global point of view, we further study the steady-state levels of αSyn* (*z*-axis) for the independent input signals of stress *S*_1_ (*x*-axis) verse the parameter *k*_5_ (*y*-axis) (Fig 5A) or the parameter *k*_6_ (*y*-axis) (Fig 5B), where the initial values of αSyn* are taken as 0, 5 and 10, respectively. The surfaces of steady-state level of αSyn* are presented with color bars and projected onto (*S*_1_,*k*_5_)- and (*S*_1_,*k*_6_)-planes as contour maps, respectively.

**FIG. 5.**
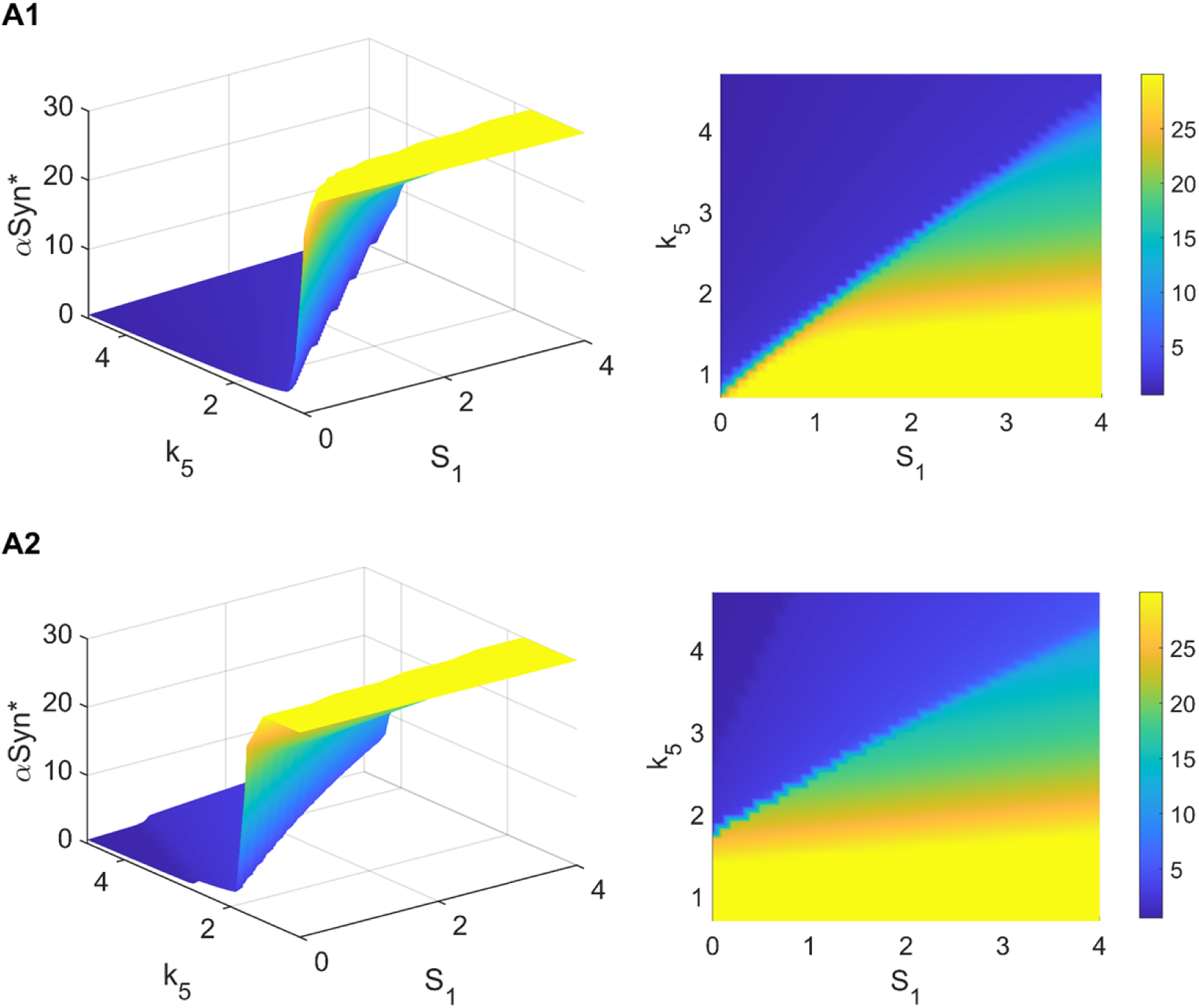
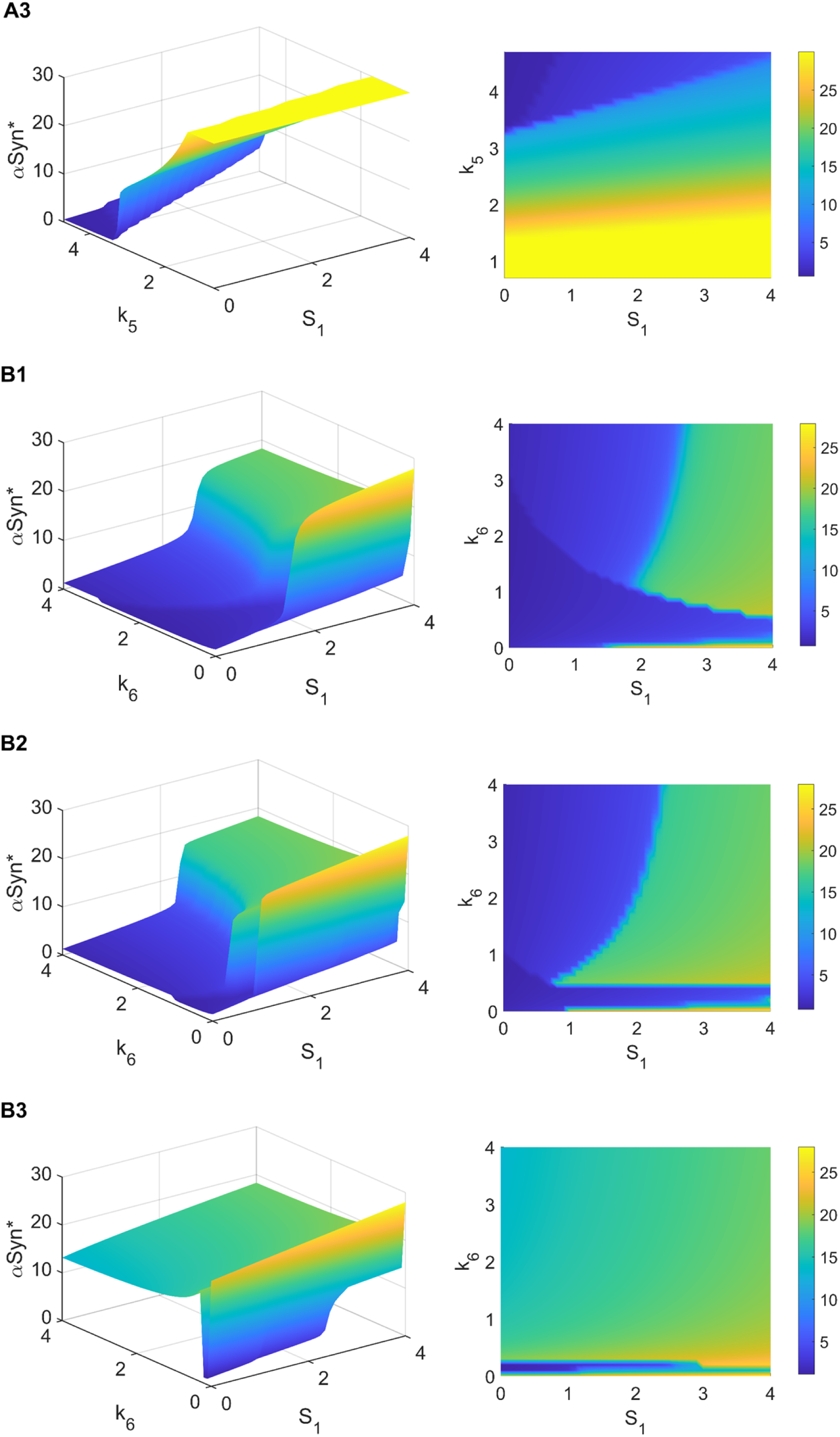
The concentration of αSyn* stays at three steady states as a function of *S*_1_ and *k*_5_ (A) or *k*_6_ (B). From top to bottom row, the initial values of αSyn* taken as 0, 5 and 10 in sequence for *k*_5_ (or *k*_6_). Each of these subgraphs shows 3D plots and their contour maps of 3D plots on the (*S*_1_, *k*_5_)- and (*S*_1_, *k*_6_)-planes.

Firstly, it is obvious that the regions of low (or high) steady-state level of αSyn* shrink (or expand) as the initial value of αSyn* increases from 0, 5 to 10 in (*S*_1_,*k*_5_)- and (*S*_1_,*k*_6_)-planes (Fig 5A and 5B), respectively. It is of particular interest in (*S*_1_,*k*_6_)-plane is that the system originated from large initial values of αSyn* almost lies in high steady-state level of αSyn* and hardly returns to low steady-state level of αSyn* (i.e., the healthy stage) regardless of the variation of *S*_1_ (Fig 5B3).

Next, we explore how the parameters *k*_5_ and *k*_6_ regulate the steady-state level of αSyn*. From the contour maps in (*S*_1_,*k*_5_)-plane (Fig 5A), it is clear that the range of low steady-state level of αSyn* (blue) expands gradually with the increase of *k*_5_. For large enough *k*_5_, there is only low stable state level of αSyn*. Indeed, the increase of *k*_5_ change the bistable switch from irreversibility to reversibility, and so the αSyn* concentration drops down to a very low level as shown in the bifurcation diagrams (Fig 3). Hence, it may be instructive to keep the parameter *k*_5_ large enough for the prevention of PD.

For very small *k*_6_, the system always especially stays in low steady-state level (blue) (Fig 3B1 and B2). Beyond a threshold value of *k*_6_, there is a sharp increase of the steady-state level of αSyn* (green) for the system initiated from a large value of αSyn* (Fig 5B3). This is due to the irreversibility of the bistable switch between the upper and lower stable states. Thus, strictly confining the parameter *k*_6_ in a very small scope may effectively prevent the occurrence of PD.

### Robustness of tristability for αSyn* to perturbation of parameters

To assess the robustness of tri-stability, we perform sensitivity analysis to detect some parameters with great influence on the tri-stable dynamic behavior. Based on the method of sensitivity analysis introduced by De Mot *et al*. [49], we figure out the extents of each parameter guaranteeing the existence of tri-stability by means of analyzing the bifurcation diagrams of the parameters (Fig 6). For example, the control parameter *k*_5_ ∈ (0.461, 4.547) for the tri-stability, that is, the value of *k*_5_ is reduced by a maximum of 82.9% or increased by a maximum of 68.4% at its default value 2.7. When tri-stability can be maintained in this range, the coexistence of tri-stability behavior is a rather robust phenomenon for the parameter *k*_5_.

**FIG. 6.**
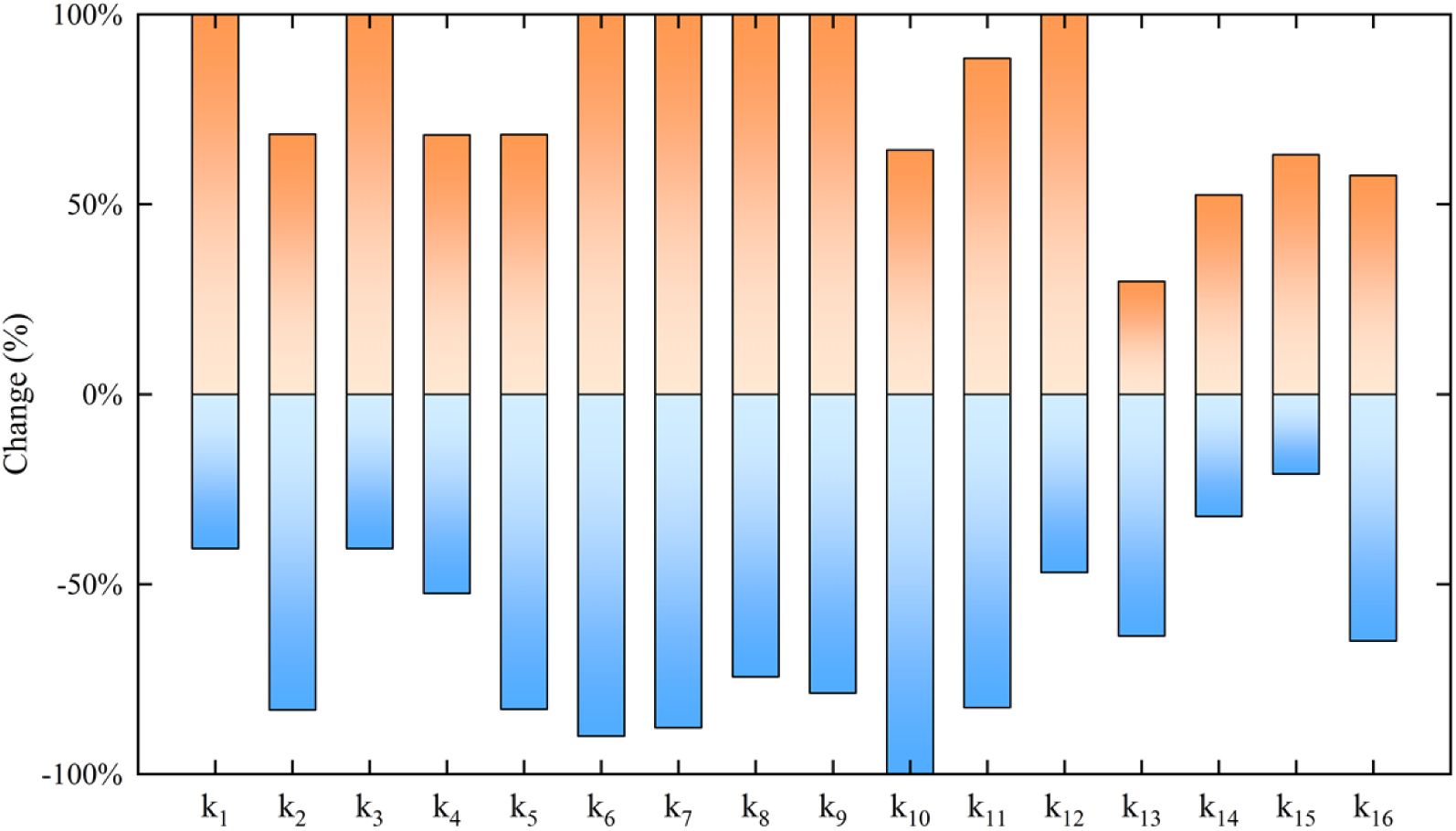
Sensitivity analysis of tri-stability. Parameters of *k*_1_ to *k*_16_ are listed, the bar denotes the range of relative variation, i.e., the percentage change in a parameter value corresponding to its default value in which tri-stability is maintained. The meaning and value of the parameters are listed in Table 1.

The above results of sensitivity analysis show that the variations of the parameters may significantly contribute to the characteristic of tri-stability for the system. Here, both Caspases-dependent rate of Beclin1 inactivation (*k*_12_) and Beclin1-dependent rate of Caspases inactivation (*k*_16_) are the key parameters involved in the double negative feedback of autophagy and apoptosis. Hence, *k*_12_ and *k*_16_ are chosen to fluctuate down and up 50 percent around their default values to evaluate the robustness of the tri-stable behaviors, by the codimension-two bifurcation diagrams in (*S*_1_,*k*_5_)- and (*S*_1_,*k*_6_)-planes (Fig 7).

**FIG. 7.**
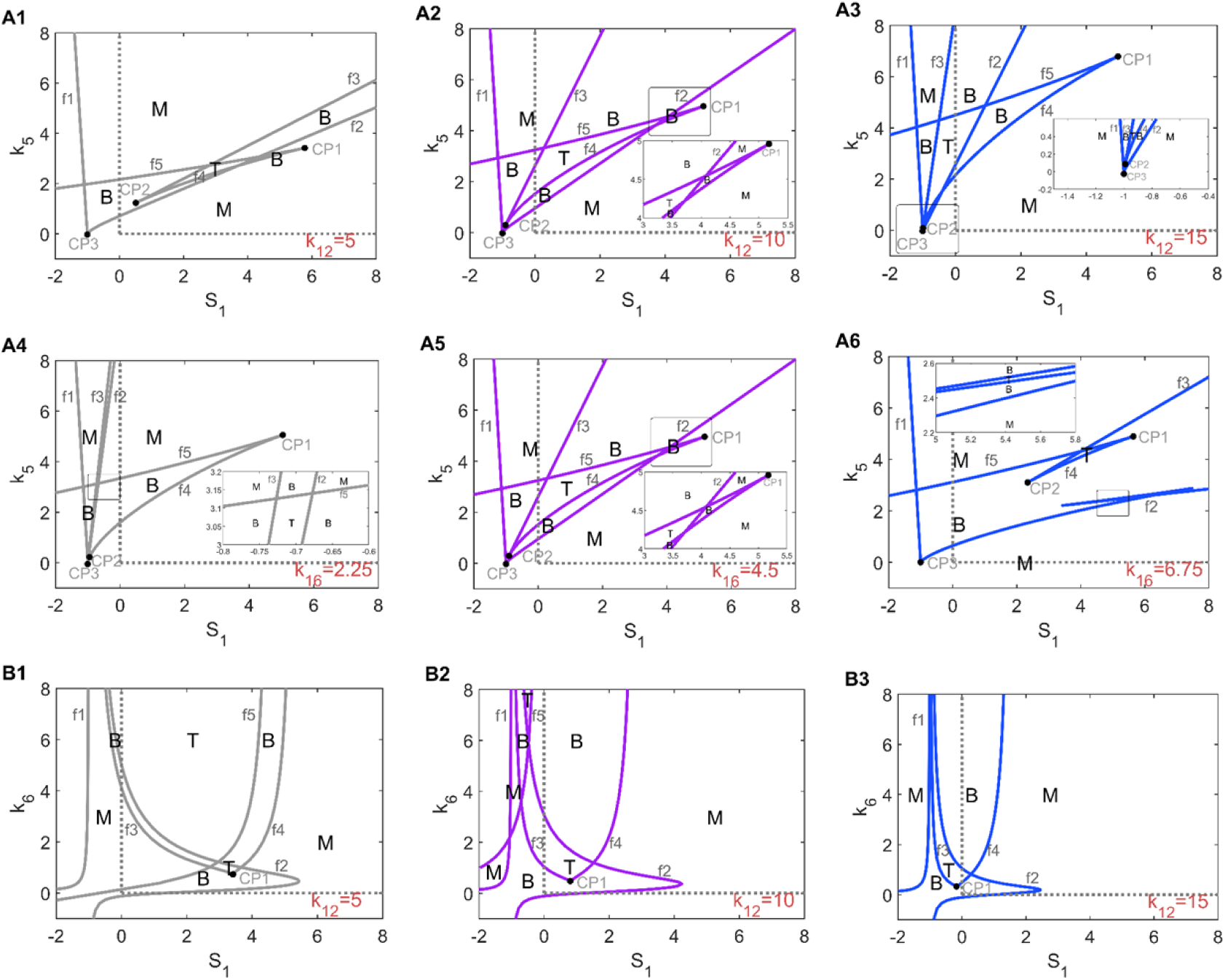
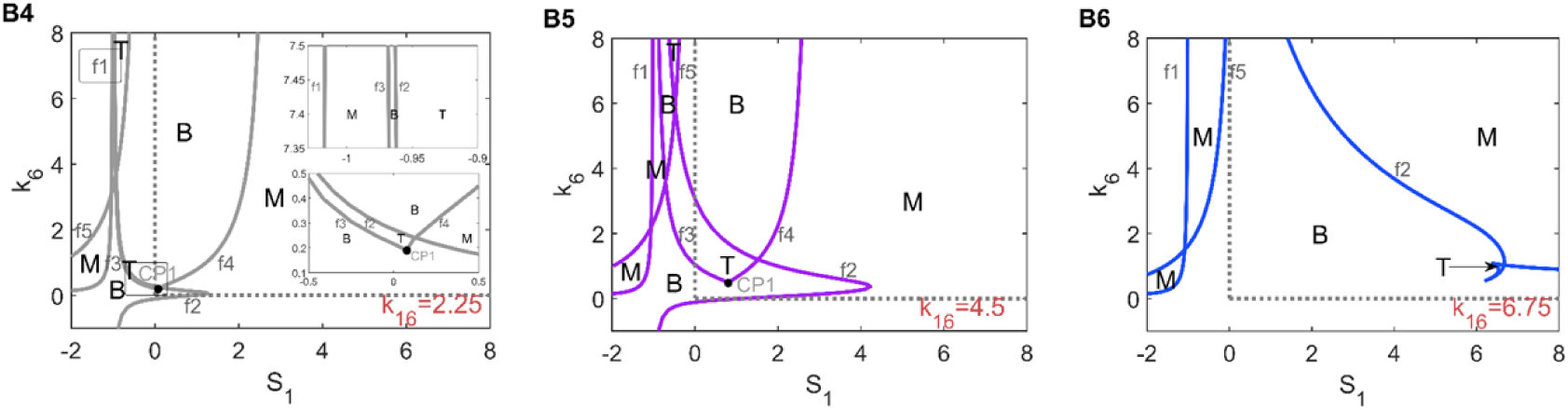
Codimension-two bifurcation diagrams of *S*_1_ verse *k*_5_ (A) or *k*_6_ (B). Parameters of *k*_12_ and *k*_16_ are chosen to fluctuate down (the third column) and up (the first column) 50 percent around their default values (the second column), respectively. T, B and M denote the regions of tri-stability, bistability and mono-stability, respectively.

We focus on the variations of the codimension-two bifurcation diagrams in (*S*_1_,*k*_5_)-plane under the perturbations of the parameters *k*_12_ (Fig 7A1-A3) and *k*_16_ (Fig 7A4-A6). Accompanied with *k*_12_ varying from small to large, the fold bifurcation curves *f*_2_, *f*_3_, *f*_4_ and *f*_5_ move gradually from right to left and the areas of the tri-stable regions T change from small to large as well. However, when the value of the parameter *k*_16_ decreases (Fig 7A5-A4), the fold bifurcation curves *f*_2_ and *f*_3_ become very close to each other; when the value of the parameter *k*_16_ increases (Fig 7A5-A6), the bifurcation curves *f*_3_ and *f*_4_ also become very close to each other. Hence, these make the tri-stable region T much narrow. All of the above indicate that the robustness of tri-stability in (*S*_1_,*k*_5_)-plane to the perturbation of *k*_12_ is better than that of *k*_16_.

We survey the disturbances of the parameters *k*_12_ and *k*_16_ to the codimension-two bifurcation diagrams in (*S*_1_,*k*_6_)-plane. Notice that the tri-stable region T changes slightly as the parameter *k*_12_ increases, even though the fold bifurcation curves *f*_2_, *f*_3_, *f*_4_ and *f*_5_ move substantially from right to left and even *f*_5_ disappears in the (*S*_1_,*k*_6_)-plane (Fig 7B1-B3). Nevertheless, the fold bifurcation curves *f*_2_ moves a lot to the left and almost collides with *f*_3_ as the parameter *k*_16_ decreases, and the fold bifurcation curves *f*_3_ and *f*_4_ vanish as the parameter *k*_16_ increases. It is seen that the area of the triable region T expands largely as the parameter *k*_16_ increases (Fig 7B4-B6). Therefore, we conclude that tri-stability in (*S*_1_,*k*_6_)-plane is more robust to the parameter *k*_12_ than to *k*_16_.

## 4 Discussion

Cloutier *et al*. [17, 18] have revealed the double positive feedback between αSyn* and ROS is essential to generate bistability while the double negative feedback between autophagy and apoptosis is associated with homeostasis. And the latest researches [50, 51] show that the existence of the critical state in multi-stability systems is an important interim for neurodegenerative disease. In our work, based on the former experimental results and theoretical foundations, we construct a new mathematic model for abnormal aggregated αSyn* degraded by autophagy-lysosome pathway.

The dynamic nature of tri-stability under the input stress *S*_1_ presents the coexistence of the lower, medium and upper stable steady states of αSyn* corresponding to the healthy, critical and diseased stages of PD, respectively. As the pathological process of PD, the critical stage can be regarded as a tipping point (or pre-disease stage) of early-warning signs for the prevention of PD [50, 51]. Under this circumstance, the existence of the critical stage acts as a barrier to sharply hamper transitions from the healthy stage to the diseased one.

Furthermore, we excavate the tri-stability of αSyn* under the regulation of Beclin1-dependent rate of αSyn* degradation (*k*_5_) and αSyn*-dependent rate of ERS activation (*k*_6_) connecting two modules of aggregation and autophagy. Increasing *k*_5_ makes the bistable switch from irreversible to reversible and so the reversibility of the bistable switch between the medium and upper stable states can pull the high level of αSyn* down for large enough *k*_5_, while the irreversibility of the bistable switch between the medium and upper stable states might hardly change the high level of αSyn* even for rather small *k*_6_

Also, we dig out a rather robust phenomenon of tri-stability dynamics by sensitivity analysis, and further focus on the robustness in the tri-stable regions in (*S*_1_, *k*_5_)- and (*S*_1_, *k*_6_)-planes with respect to the Caspases-dependent rate of Beclin1 inactivation (*k*_12_) and the Beclin1-dependent rate of Caspases inactivation (*k*_16_). The results illustrate that autophagy significantly contributes to the clearance of aggregated αSyn* during the treatment process of PD from the dynamic point of view [33, 41].

Our model built the connection of proteins clearance mechanism of autophagy in degrading aggregated αSyn*, and puts the experimental and theoretical frameworks together for understanding the precise regulation of autophagy to degrade αSyn*. The medium stable steady state in this model would create a barrier to avoid the system to easily transit from the lower stable steady state to the upper one, so it is better to let neuron cells stay at this critical stage for appropriate treatments to prevent more aggregation of αSyn* causing PD worse. Therefore, our work may provide meaningful outcomes and new insights to the prevention or treatment of PD. Moreover, the advantage of finding tri-stability is important for further dynamic modelling of PD pathological networks. The crucial functions of the medium state controlled by more related molecular should be considered in the mathematical modelling and dynamic analysis of PD in future. The important properties of tri-stability may also be beneficial to future experimental studies of PD.

## Data Availability Statement

All relevant data are within the manuscript.

## Acknowledgments

The authors would like to thank Prof. Qishao Lu in Beihang University for insightful discussions and valuable comments for this work.

## Author Contributions

**Conceptualization:** Bojie Yang, Zhuoqin Yang

**Formal analysis:** Bojie Yang, Zhuoqin Yang, Heng Liu

**Methodology:** Zhuoqin Yang

**Software:** Bojie Yang, Heng Liu

**Supervision:** Zhuoqin Yang

**Writing – original draft:** Bojie Yang

**Writing – review & editing:** Zhuoqin Yang

